# Optimized DNA isolation method for microbiome analysis of human tissues

**DOI:** 10.1101/2020.08.25.267641

**Authors:** Carlijn Bruggeling, Daniel R. Garza, Soumia Achouiti, Wouter Mes, Bas E. Dutilh, Annemarie Boleij

**Affiliations:** Department of Pathology, Radboud Institute for Molecular Life Sciences (RIMLS), Radboud university medical center (Radboudumc), the Netherlands; Center for Molecular and Biomolecular Informatics (CMBI), Radboud Institute for Molecular Life Sciences (RIMLS), Radboud university medical center (Radboudumc), the Netherlands; Department of Animal Ecology & Physiology and Department of Microbiology, Institute for Water and Wetland Research (IWWR), Radboud university, the Netherlands; Theoretical Biology and Bioinformatics, Science for Life, Utrecht University, Utrecht, The Netherlands

## Abstract

Recent advances in microbiome sequencing have rendered new insights into the role of the microbiome in human health with potential clinical implications. Unfortunately, developments in the field of tissue microbiomes have been hampered by the presence of host DNA in isolates which interferes with the analysis of the bacterial content. Here, we present a DNA isolation protocol from tissue samples including reduction of host DNA without distortion of microbial abundance profiles. We evaluated which concentrations of Triton and saponin lyse host cells and leave bacterial cells intact, which was combined with DNAse treatment to deplete released host DNA. We applied our protocol to extract microbial DNA from *ex vivo* and *in vivo* acquired human colon biopsies (∼2-5 mm in size) and assessed the relative abundance of bacterial and human DNA by qPCR. Saponin at a concentration of 0.0125% in PBS lysed host cells, resulting in a 4.5-fold enrichment of bacterial DNA while preserving the relative abundance of *Firmicutes, Bacteroidetes, γ-Proteobacteria* and *Actinobacteria*. Our protocol combined with shotgun metagenomic sequencing revealed a colon tissue microbiome profile with a Shannon diversity index of 3.2 and an UniFrac distance of 0.54, which is comparable to reported numbers based on amplicon sequencing. Hereby, we present the first protocol for enriching bacterial DNA from tissue biopsies that allows efficient isolation of rigid Gram-positive bacteria without depleting the more sensitive Gram-negative bacteria. Our protocol facilitates analysis of a wide spectrum of bacteria of clinical tissue samples improving their applicability for microbiome research.

## Introduction

The rapidly growing field of microbiome research is steadily revealing a role of the microbiome in human health and diseases. Functions of the gut microbiome are diverse and essential for many biological processes involved in metabolism, tissue homeostasis and immunity (1). Changes in microbiome composition have been associated with a wide variety of diseases, ranging from intestinal inflammatory diseases to colorectal cancer to diseases outside the gastrointestinal tract (1). Such compositional changes are well-studied by microbiome profiling through sequencing of DNA isolates. While a vast amount of research has been performed on stool, recent technologies have facilitated bacterial profiling on colon tissues, which allows more localized analysis (2) and may be more accurate in differentiating between healthy and diseased states (3). Importantly, DNA isolation methods have a major impact on the evaluation of microbiota composition (3-11). Hence, a well-developed and standardized protocol for stool and tissues will contribute to consensus in microbiome research.

The study of microbiome composition of solid tissue samples however, does not come without challenges. Whole tissue isolates contain large bulks of host DNA, overshadowing the presence of single-cell organisms and viruses. While polymerase chain reaction (PCR) is a valuable technique to identify minority sequences, the field of microbiome research is slowly moving towards shotgun metagenomic sequencing as a preferredmethod. Shotgunmetagenomic sequencing allowsanalysis of all sequences in the DNA isolate, resulting in an increased species detection with higher accuracy (12). Another major advantage of this technique is the ability to discriminate between microbial species and analyze its gene content including potential virulence factors (12). This may be crucial to discriminate between a pathogen and a commensal bacterium at species level (13). Unfortunately, the application of shotgun metagenomic sequencing to study the microbiome of human tissue is complicated due to the large amount of human DNA: large amounts of input DNA are required to reach enough depth for sequence analysisof the microbial DNA fraction. Reduction of human DNA in tissue isolates is required to increase sensitivity of shotgun metagenomic sequencing microbiome analysis of tissue.

Various methods have been developed to improve the bacterial to human DNA ratio. These methods include filtering out human cells by cell size (14), antibody-mediated filtration of human DNA by targeting non-methylated CpG dinucleotide motifs (14, 15) and human-specific cell lysis followed by DNA degradation (7, 11, 14, 15), of which the latter results in most efficient bacterial DNA enrichment (11, 14). Hence, bacterial DNA enrichment contributes to the identification of minority species and to a higher resolution of the microbial genomes present in the sample, rendering improved bacterial classification and analysis of genes of interest.

One of the caveats of bacterial DNA enrichment is that the method of DNA isolation affects the microbiome profile (7, 9, 11, 14-17). Bacteria differ in their susceptibility to lysis, resulting in the tendency of some bacteria to lyse too early during the isolation method (15, 16), while other bacteria may require extra steps to release their DNA, e.g. by mechanical lysis through bead-beating (10, 18). Addition of mechanical lysis has improved isolation of Gram-positive bacteria (4, 9, 16), without impairing the isolation of Gram-negative bacteria (19). Additionally, enzymatic lysis with mutanolysin may help to identify more Gram-positive bacteria (4, 20). The ultimate goal is to increase the bacterial to human DNA ratio and have a DNA isolate that closely reflects the bacterial composition of the sample.

The immense improvement by shotgun metagenomic sequencing in the field of the microbiome has been based on clinical stool samples; not tissue. Thereby,the study of the bacteria that reside in closest proximity to the host are left outside consideration, along with crucial information about their localization in the gut (e.g. colonic segment or localization to tumors). Current protocols can be optimized for analysis of the tissue microbiome, for which we present our improved method in this paper. Our method combines important elements of the currently best performing methods for DNA isolation so far: bacterial DNA enrichment, mutanolysin treatment, heat-shock and bead-beating. Our protocol is designed for an unbiased isolation of diverse microbes rendering efficient lysis of Gram-positive bacteria, while maintaining efficient isolation of Gram-negative bacteria. The inclusion of our fine-tuned microbial DNA enrichment strategy enriches the bacterial content and results in a reproducible analysis of microbial profiles of biopsies ranging from ∼2-5 mm. Thus, this method will contribute to reproducible research in the field of microbiome composition and functionality and will be of value not only for gut-related tissue microbe analysis, but also for those tissues where microbes are underrepresented (e.g. fish gills).

## Methods

### Collection of human colon biopsies

*Ex vivo* residual resected colon material was obtained at the department of pathology of the Radboudumc in Nijmegen between 2017 and 2018, in accordance with Dutch legislation. No approval from a research ethics committee was required for the study of residual colon resections, because anonymous use of redundant tissue for research purposes is part of the standard treatment agreement with patients in the Radboudumc, to which patients may opt out. None of the included patients submitted an objection against use of residual materials and all material was processed anonymously. Biopsies were resected with a scalpel, resulting in biopsies up to an estimated size of 5 mm. Alternatively, a biopsy forceps was used to make biopsies of about 2 mm that were used as a proxy for biopsies taken during colonoscopy. After collection, biopsies were snap-frozen in cryo-tubes in liquid nitrogen and stored at −80°C.

*In vivo* collected forceps biopsies for shotgun metagenomic sequencing were obtained from patients that came for a screening colonoscopy and participated in either of the two studies: the BBC study (NL57875.091.16), which were solely genetically confirmed Lynch syndrome patients, the BaCo study (NL55930.091.16) which included ulcerative colitis patients and patients without known colon diseases. Samples were collected between 2017 and 2018 in Radboudumc Nijmegen. Both studies were approved by the Internal Revenue Board CMO-Arnhem Nijmegen (CMO 2016-2616 and CMO 2016-2818) and the board of the Radboudumc. Patients whom had taken antibiotics within the last 3 months prior to the colonoscopy were excluded. All patients were older than 18 years and signed an informed consent. Biopsies were snap-frozen in cryo-tubes in liquid nitrogen instantly after collection and stored at −80°C.

### Bacterial DNA isolation protocol

The bacterial DNA isolation strategy involved bacterial DNA enrichment through human cell lysis and DNAse treatment (see figure 1, upper part), which was followed up by our previously optimized bead-beating protocol (see figure 1, lower part) (21). Whereas the bead-beating protocol remained unchanged throughout this paper, two alternative strategies were tested for the bacterial DNA enrichment. For the first strategy, the Molzym DNA isolation (Ultra-Deep Microbiome prep, Molzym) kit was used. The manufacturer’s protocol was followed until and including the molDNAseinactivation step. Subsequently, the bead-beating protocol was applied to assist in mechanical bacterial cell lysis, because this was shown to result in a higher bacterial signal in qPCR (supplementary figure 1). For the second strategy, we established our own alternative protocol including proteinase K (19133, Qiagen) for tissue digestion, Phosphate buffered saline (PBS) (Braun, 220/12257974/1110) containing saponin (47036-50G, Sigma-Aldrich) or Triton for selective lysis, and TurboDNAse (AM2239, Qiagen) for host DNA removal. We evaluated the effect of Triton or saponin at different concentrations forhuman cells and experimented what was the best moment to include the biopsy wash (point A or B) in the DNA-isolation process (Figure 1).

**Figure 1.**
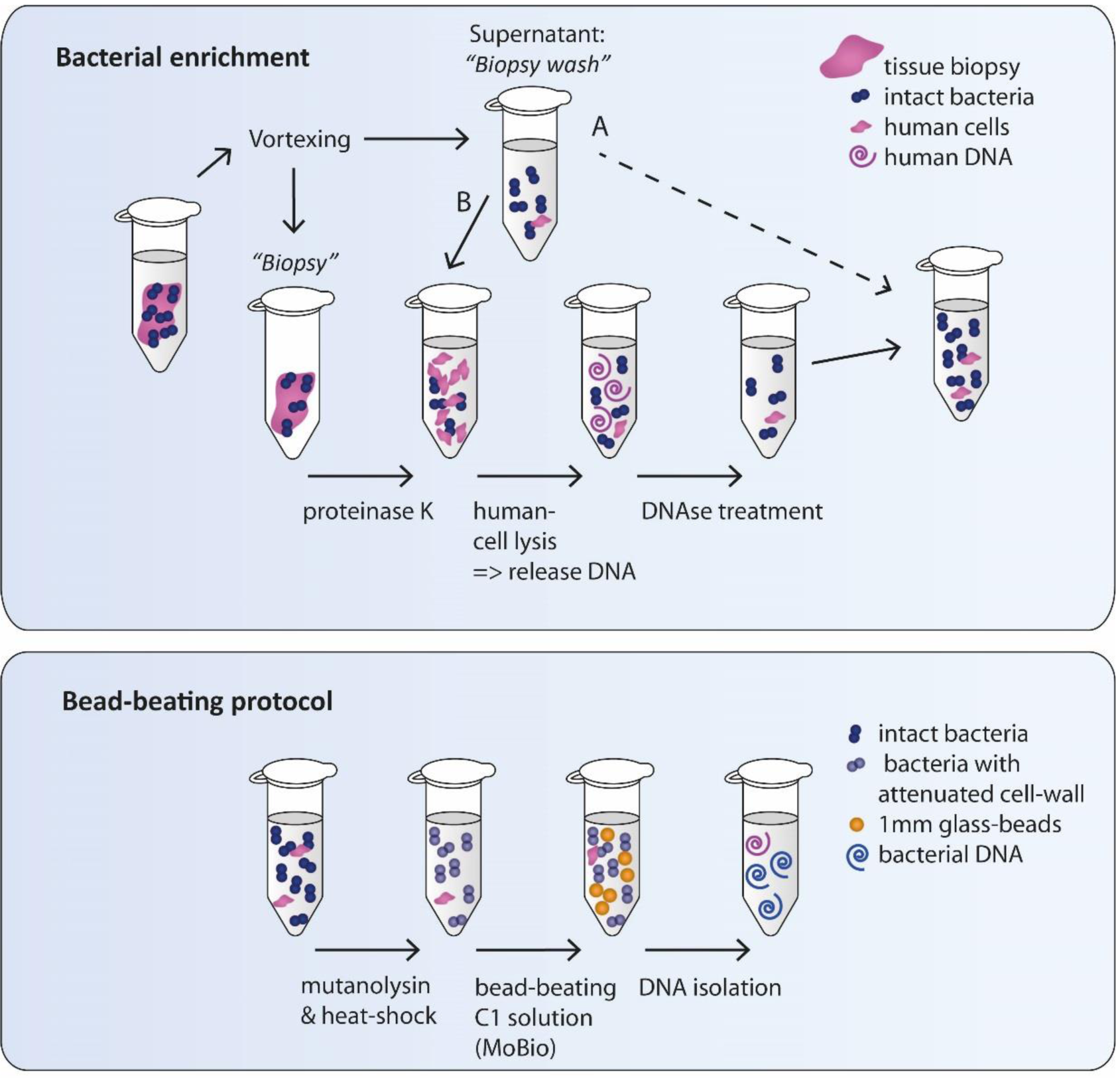
Schematic drawing of DNA isolation protocol strategy 2. <u>A. Bacterial enrichment:</u> A tissue biopsy is vortexed in PBS to separate bacteria from the biopsy. The biopsy is retrieved for digestion with proteinase K, while the supernatant (biopsy wash) is saved on ice and added back for DNA isolation at a later timepoint (timepoint A or B; B in the final protocol). Bacteria in the biopsy wash are thereby minimally exposed to reagents that could cause possible lysis. Subsequently, 0.0125% saponin in PBS is added to the cell suspension inducing lysis of human cells, but not bacterial cells. DNA in the supernatant is depleted through DNAse treatment. The remaining sample has reduced human DNA content and still intact bacteria. <u>B. Bead-beating protocol:</u> The sample is further processed by our previously optimized bead-beating protocol. Mutanolysin treatment followed by heat-shock are applied to attenuate cell -walls of Gram-positive bacteria (e.g. *Streptococci* and *Actinobacteria*) to make them more susceptible for mechanical lysis. Subsequently, the sample is bead-beated with 1 mm glass-beads in C1 buffer of the Powerlyser powersoil DNA isolation kit and further isolated according to the manufacturer’s protocol. The resulting DNA isolate is enriched for bacteri al DNA.

The lysis of bacterial cells included treatment with 0.5 KU/mL mutanolysin (SAE0092, Sigma Aldrich), heat-shock and buffer C1 of the DNAeasy powerlyzer Powersoil kit from Qiagen (previously known as the MoBio Powerlyzer PowerSoil DNA isolation kit from MoBio). Bead-beating was performed in the Magnalyser (Roche) at 6400 rpm for 20 seconds twice, with 30 seconds on ice in between. After bacterial lysis the manual of the DNA-isolation kit was followed. The final protocol is provided in supplementary file 1. Our final bacterial enrichment protocol (figure 1, upper part and supplementary file 1) was also tested by an independent laboratory (Institute for Water and Wetland Research, Radboud University) for isolation of bacteria from zebrafish gills, but in combination with CTAB extraction instead of the MoBio DNA isolation kit (supplementary file 2).

### Bacterial culturing

*Collinsella intestinalis* (DSM13280), *Bacteroides vulgatus* (3775 SL(B)10), *Escherichia coli* (NTB5) and *Streptococcus gallolyticus* subsp. *gallolyticus* (UCN34) were cultured on Brain-Heart-Infusion agar plates supplemented with yeast extract L-cysteine Vitamin K, and Hemin (BHI-S; ATCC medium 1293). *C. intestinalis* and *B. vulgatus* were grown on plates for 48 hours under anaerobic conditions before transfer to liquid medium for 48-72 hours at 37°C. *E. coli* and *S. gallolyticus* were grown overnight on plated under aerobic conditionsbefore transfer to liquidculturingin BHI for 24 hours at 37°C. Bacteria were pelletedby centrifugationat 4600 rpm for 10 minutes and frozen at −20°C. Bacterial pellets were thawed and dissolved in PBS until 1 optical density (OD at 620 nm) of which 50 µL was used for experiments to determine bacterial DNA release by Triton and saponin treatment.

To create a mock community, 1 OD bacterial PBS suspensions were mixed in 400 µL (40% *B. vulgatus*, 30% *E. coli*, 20%, *S. gallolyticus* and 10% *C. intestinalis*) and were pelleted for each experimental condition.

### Bacterial DNA release by treatment with Triton and saponin

Bacteria were dissolved in PBS with final concentrations of Triton or saponin of 0.1%, 0.025%, 0.0125% and 0.006%. Bacteria were incubated for 30 minutes at 37°C with a soap or PBS only. Samples were centrifuged at 10000 x g for 10 minutes and the DNA concentration was measured with Qubit Fluorometer 2.0 (Thermofisher scientific) using the Qubit dsDNA HS assay kit (Q32856, Thermofisher). A Mann-Whitney U-test was used to compare the DNA in the supernatants of samples exposed to a soap versus PBS.

### Effects of Saponin 0.0125% on human tissue lysis

To test whether saponin 0.0125% was able to induce human cell lysis, resected human colon biopsies of an estimated size of 5 mm were processed according to our optimized protocol up to the step of selective cell lysis using saponin (see figure 1 and supplementaryfile 1). During this last step, cell pellets were incubated with either 0.0125% saponin or PBS in turboDNAse buffer, but without turboDNAse enzyme. Samples were incubated at 37°C for 30 minutes to lyse the cells and the supernatant was cleared from cell debris by two centrifugation cycles of 10 minutes at 10000 x g at 4°C. DNA in the supernatant was precipitated with 100% ethanol and centrifuged at 10000 x g at 4°C for 20 minutes. Precipitated DNA was washed with 70% ethanol and centrifuged at 10000 x g at 4°C for 20 minutes. Lastly, DNA was air dried and resuspended dH_2_0.

### Quantitative Real-time PCRs for 16s rRNA

Each reaction for qPCR consisted of 0.4 µM forward primer, 0.4 µM reverse primer, 1X Power SYBR Green (A4368702, Applied biosystems). The amount of DNA in each reaction was 1 ng and 0.1 ng for biopsies that were ∼5 mm and ∼2 mm, respectively. Primers for host (human or zebrafish) and bacteria (all bacteria, *Firmicutes, Bacteroidetes, γ-Proteobacteria* and *Actinobacteria*) were used and evaluated before (21-23) and are reported in our Supplementary table 1 (22-27). qPCRs were performed with a 7500 Fast Real-Time PCR system (Applied Biosystems®). Samples were heated to 50°C for 2 minutes, 95°C for 10 minutes, 30 cycles of 95°C for 15 seconds and 60°C for 1 minute, followed by a continuous sequence of 95°C for 15 seconds, 60°C for 1 minute, 95°C for 30 seconds and 60°C for 15 seconds. Melting curves were generated to evaluate the specificity of the PCR-product.

DNA isolated from the mock community (described above) was used as a positive control. Only for supplementary figure 1, a human fecal isolate was used as a positive control. DNA isolated from human blood served as a negative control.

### Statistical analysis of qPCRs

To evaluate differences in bacterial content between samples, the universal 16S rRNA signal of the sample was calibrated using the universal 16S rRNA signal of the positive control (ΔCt); a mock community isolate that resembles the gut microbiome. Fold difference was calculated by 2^∧-ΔCt^. To study bacterial composition, the 16S rRNA signal of *Firmicutes, Bacteroidetes, Actinobacteria* or *γ-Proteobacteria* was calibrated with the 16S rRNA signal of the Universal signal of the same sample (ΔCt). Subsequently, the ΔCt was compared to the ΔCt in a control sample (ΔΔCt). Fold difference was calculated by 2^∧-ΔΔCt^. Paired samples were analysed witha paired-T test. In case of unmatched samples, the Mann-Whitney U-test was used for comparison.

A Friedman test was used to evaluate which soap resulted in the most similar bacterial composition to PBS. All statistical tests were performed using Graphpad Prism version 5.0.

### Shotgun metagenomic sequencing of human colon biopsies

DNA was isolated using the DNeasy Powerlyzer Powersoil kit (Qiagen), as described in supplementary file 1. DNA concentration was measured as described previously 521 human colon tissue DNA isolates were send to Novogene Bioinformatics Technology Co., Ltd in Hongkong for sequencing. Samples were processed using low input NEBnext library preparation and paired-end sequencing was performed on the Illumina Novaseq 6000 with 350 bp insert size and a read length of 150 bp. 1.2 GB output data in FastQ format was guaranteed per sample. Samples were measured for DNA concentration, construct length and a quality check was performed on the library preparation. 13 samples were not sequenced due to failed library preparation.

### Bioinformatics analysis

Quality control, trimming, and removal of adaptors was performed using FastQC version 0.11.9 and trimmomatic version 0.35. An assembly dataset was generated by filtering out the human reads using BBMap version 38.84 with the GRCh38 version of the human genome. Filtered reads were assembled with metaSPAdes version 3.13.1. The taxonomic classification of contigs was determined with CAT v. 4.6 (PMID:31640809) using the NCBI NR as database for taxonomic assignments. bwa version 0.7.17 and samtools version 1.9 were used to map all the reads to the classified contigs and the human genome and to estimate the coverage statistics. Only samples with more than 2.0e04 bacterial reads were used, resulting in 225 metagenomes derived from human colon biopsies with an average of 11 million reads per sample. Shannon diversity (alpha) and the UniFrac diversity (beta)(28) were estimated from the taxonomic distribution of reads at the genus level. Diversity indices and phylum-level classifications were compared to values obtained from literature (29-32)

## Results

### Whole tissue digestion including PBS wash is required to capture the collective tissue microbiome

It is hypothesized that a major bulk of human DNA in the microbial DNA isolate could be avoided by only isolating DNA from washed tissue (biopsy wash). To test this, the biopsy and biopsy wash were isolated separately with the Ultra-Deep Microbiome prep-kit (Molzym) in combination with our bead-beating protocol. While biopsies were isolated with the full protocol including tissue digestion, selective lysis and removal of human DNA using strategy 1 (see methods), these steps were omitted for the biopsy wash (path A in Figure 1). Similar universal bacterial 16S rRNA signals were obtained from DNA isolates of the biopsy wash and biopsies (Figure 2). This suggests that isolating DNA from the biopsy wash would only represent a selective part of the microbial community and hence isolation of the whole biopsy includingthe biopsy washis necessary to capture the collective tissue microbiome.

**Figure 2.**
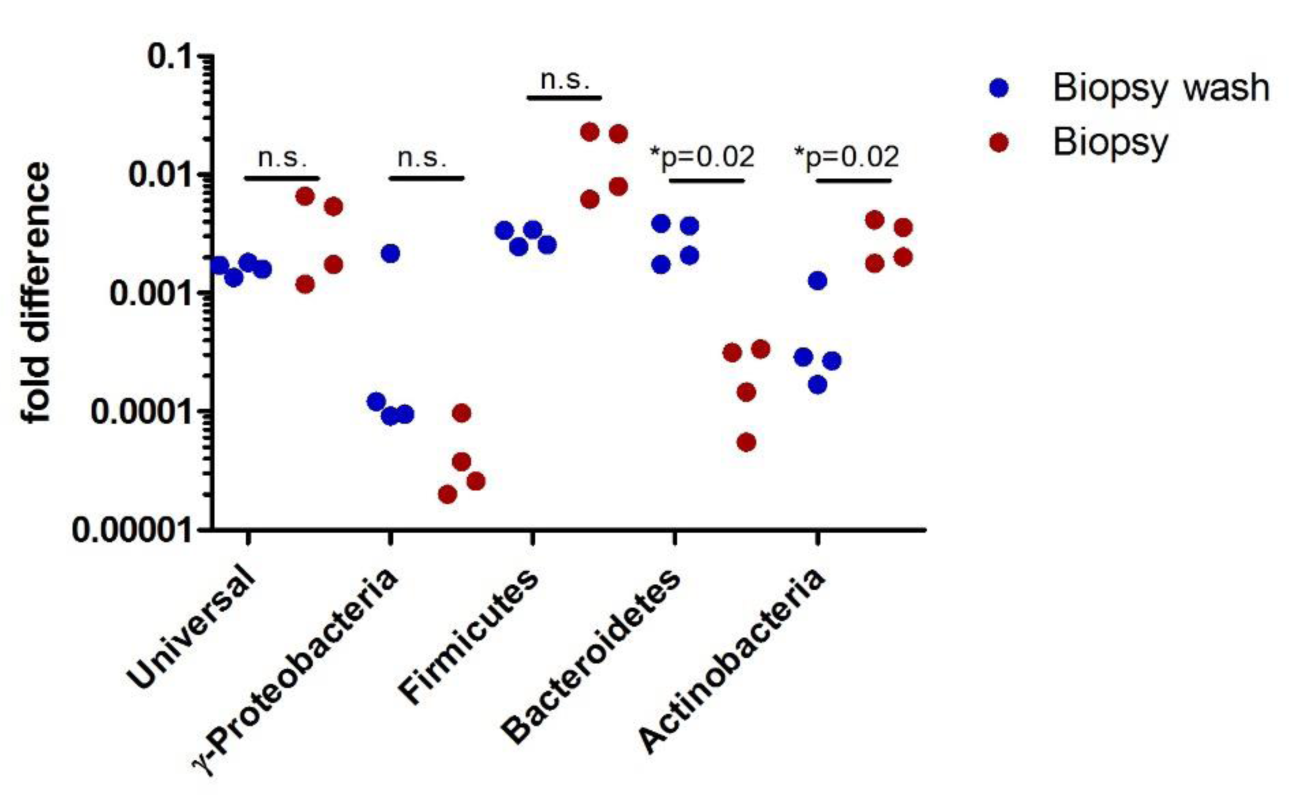
Whole tissue digestion is required to isolate all bacteria. Two matched biopsies (∼5 mm) were washed in PBS, after which DNA of the Biopsy wash and the Biopsy was isolated separately. For every DNA isolate a duplicate was run, of which each value is plotted relative to the mock community (ΔCt). Paired T-tests revealed that DNA from the biopsy isolates contained a similar bacterial fraction, albeit with fewer *Bacteroidetes* and more *Actinobacteria*. Hence, whole tissue digestion is required to analyze the complete bacterial component of the tissue.

### DNA-isolation using strategy 1 changes microbial composition

Interestingly, the biopsy wash appeared to have relatively more Gram-positive and fewer Gram-negative bacteria compared to the microbiota remaining in the matched biopsy. This difference was significant for *Bacteroidetes* (p=0.02) and *Actinobacteria* (p=0.02) (figure 2). Theoretically, this discrepancy could be caused by isolation of different bacterial populations: e.g. bacteria in the outer mucus layer (biopsy wash) and inner mucus layer or within the tissue (biopsy) of which the latter may remain attached to the biopsy after vortexing in PBS. Alternatively, we hypothesized that one of the buffers in the Ultra-deep microbiome prepkit could cause premature lysis of especially Gram-negative bacteria to which the biopsy washes were not exposed. Therefore, we tested the effect of strategy 1 on bacterial composition by applying DNA isolation on a pure bacterial culture; a mock community. We compared the full protocol (similarly to the biopsy) or a part of the protocol (similarly to the biopsy wash, Path A in Figure 1). We found that the full strategy 1 protocol, which includes selective cell lysis and DNAse treatment, resulted on average in a 15-fold lower signal of γ-Proteobacteria (p=0.03) and a 27-fold lower signal of *Bacteroidetes* (p=0.03) as opposed to the incomplete protocol (see supplementary figure 2). This suggests that strategy 1 disfavors isolation of Gram-negative bacteria versus Gram-positive bacteria.

### The microbial community composition is preserved with 0.0125% saponin while selectively lysing human cells

Because strategy 1 changed microbial composition, strategy 2 was established using similar, but tweakable steps, including tissue digestion with proteinase K, selective humancell lysis withsoaps and DNAse treatment to remove host cell DNA after lysis. First, we tested which soap would effectivelylyse human cells without affecting the composition of the microbiome. Hence, we tested whether treatment with different concentrations of Triton and saponin would result in bacterial DNA release (eDNA).

First, pure bacterial cultures of *Streptococcus gallolyticus (Firmicutes), Bacteroides vulgatus (Bacteroidetes), Echerichia coli (γ-Proteobacteria)* and *Collinsella intestinalis (Actinobacteria)* (figure 3a) were exposed to Triton and saponin. While *C. intestinalis* was resistant to lysis under all conditions, *B. vulgatus* and *S. gallolyticus* were susceptible to lysis in the presence of Triton, with higher concentrations leading to more eDNA. Triton did not affect the amount of eDNA of *E. coli* and *C. intestinalis*. Saponin was shown to be a milder soap, as it only increased eDNA of *E. coli* at a concentration of 0.1%.

**Figure 3.**
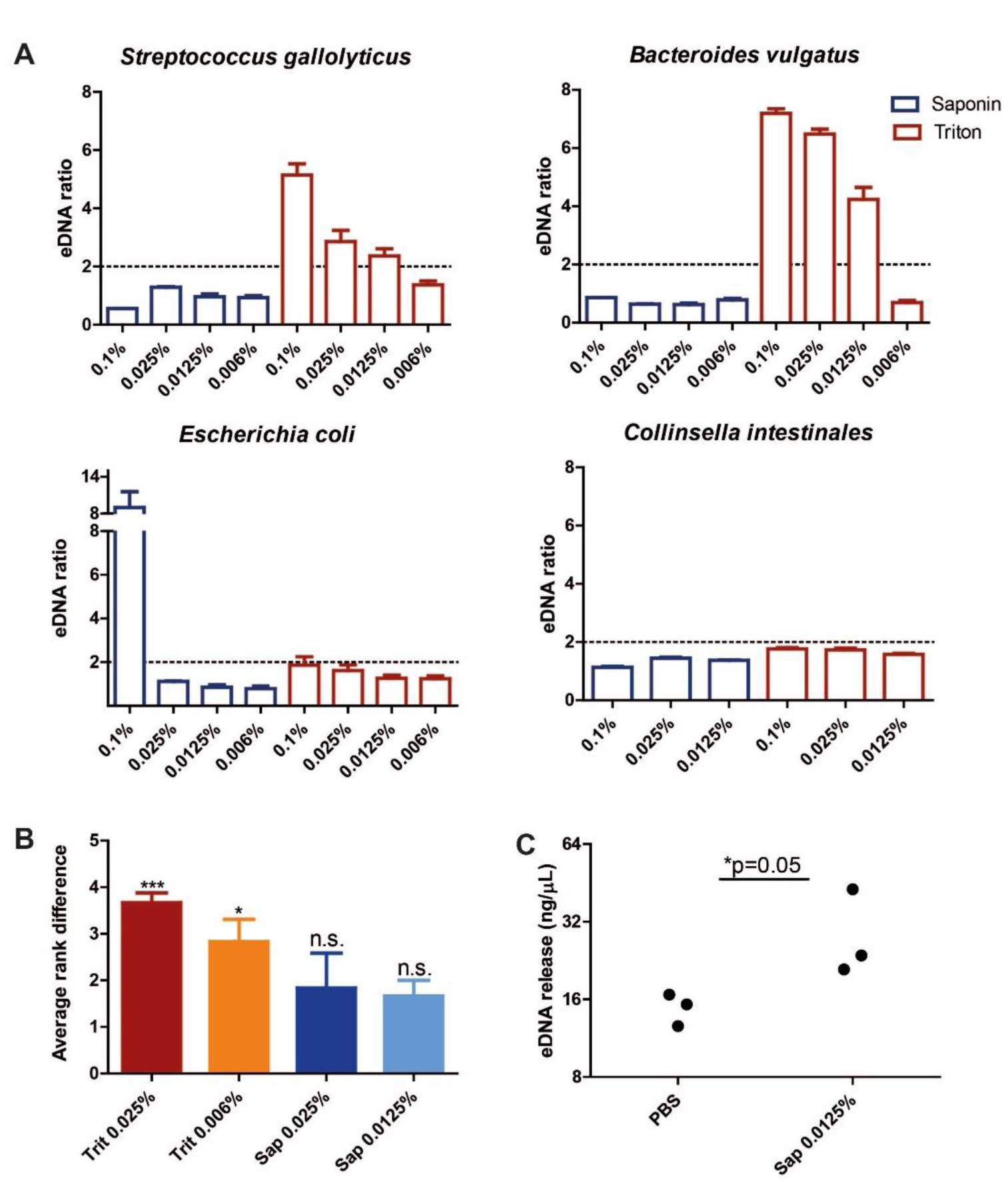
Saponin 0.0125% induces human cell lysis, without inducing bacterial cell lysis. The effect of Triton and saponin on bacterial cell lysis was measured. This experiment was performed for *Streptococcus gallolyticus (Firmicutes), Bacteroides vulgatus* (*Bacteroidetes*), *Escherichia coli* (*γ*-*Proteobacteria*) and *Collinsella intestinales* (*Actinobacteria*). An increase of more than 2 was considered relevant. Results show that Triton affects bacterial cell lysis in *Streptococcus gallolyticus* and *Bacteroides vulgatus*, but not in *Escherichia coli* and *Collinsella intestinalis*. Saponin only induced cell lysis at 0.1% in *E. coli*. B) Biopsies were isolated with strategy 2 in combination with Triton (Trit) and saponin (Sap) at different concentrations. The relative bacterial signal for *Firmicutes, Bacteroidetes, Actinobacteria* and *γ-Proteobacteria* was calibrated with the universal 16S rRNA signal (ΔCt) and was compared to PBS (ΔΔCt). Similarity to PBS was calculated through ranking using the Friedman test. Both saponin concentrations most closely resembled bacterial composition in PBS and hence preserved bacterial composition at phylum level in the colon biopsies. C) DNA release of biopsies was measured after exposure to either PBS or saponin 0.0125%. More external DNA (eDNA) was measured after incubation with saponin 0.0125% (p=0.05), suggesting that human cell lysis was induced, although eDNA was also detected in the sample with PBS alone.

Secondly, it was tested whether Triton and saponin would change the bacterial composition of tissue from 2 patients (patient 1 and patient 2). DNA was isolated using the protocol including either saponin (0.0125% or 0.025%) or Triton (0.025% or 0.006%) and the relative abundance of *Firmicutes, Bacteroidetes, Actinobacteria* and *γ-Proteobacteria* was compared to isolations performed without soap (PBS). For each phylum, the soap creating the lowest distance to PBS was ranked 1, followed by rank 2, 3, and 4 (supplementary figure 3). Saponin 0.0125% led to the smallest difference in abundance with PBS across all bacterial phyla (supplementary figure 3, figure 3b). Triton 0.006% and Triton 0.025% ranked significantly higher (p<0.05 and p<0.001 respectively) (figure 3b). Additionally, the *Firmicutes* to *Bacteroidetes* ratio was only maintained in the saponin 0.0125% condition (supplementary figure 4). Thus, saponin 0.0125% preserved relative bacterial composition within the samples.

Thirdly, we tested whether saponin 0.0125% would mediate human cell lysis by exposing 2 sets of 3 tissue homogenates (size: ∼5 mm) to either PBS or saponin 0.0125%. The tissue supernatant treated with saponin contained more than twice the amount of eDNA compared to tissue in PBS only (p=0.05) (figure 3c). This shows that exposure of tissue to saponin 0.0125% induces selective lysis of host cells, while keeping bacterial cells intact and maintaining bacterial composition.

### Strategy 2 increases the bacterial to human signal

After specific eDNA release of human tissue, DNAse treatment should be performed to degrade the released human DNA. Degradation of eDNA significantly reduced free DNA in the supernatant (figure 4b). The significant lower DNA yield after DNAse treatment was associated with an increased bacterial signal in qPCR (p=0.004) (figure 4a), which is indicative of a greater bacterial to human DNA fraction in the tissue DNA isolate.

**Figure 4.**
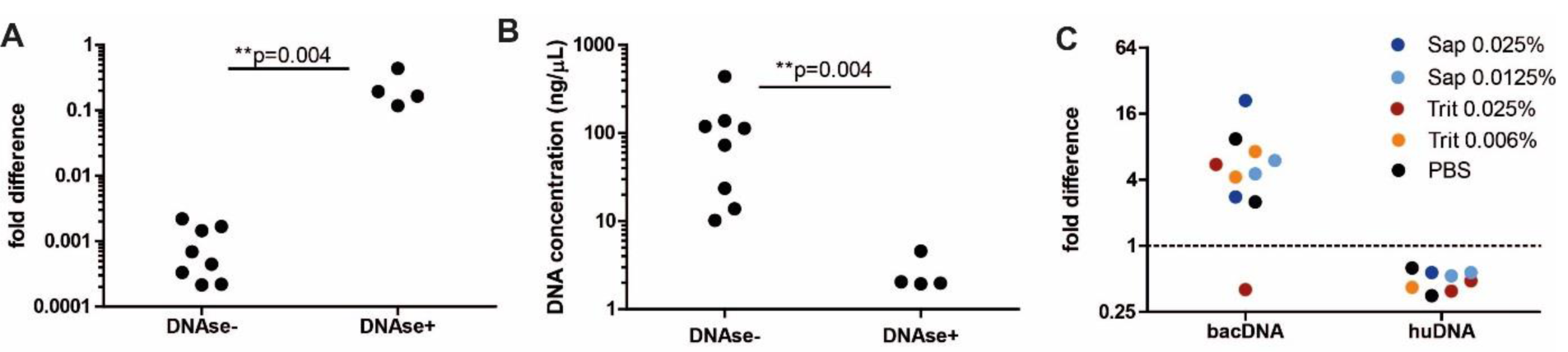
DNAse treatment lowers total DNA yield and improves bacterial to human DNA signal. A+B) To test the effectiveness of bacterial DNA enrichment, isolations were performed on tissues (∼5 mm) with or without the biopsy wash included in the DNAse treatment (DNAse+ and DNAse-respectively, which represent path B and A respectively in figure 1). DNAse treatment results higher bacterial signal (p=0.004) (A) which corresponds with a lower DNA yield (p=0.004) (B). These results suggest that DNAse treatment on the PBS wash enriches the bacterial DNA content of the isolate, illustrating that PBS wash should be included during DNAse treatment (path B in figure 1). C) To test the effect of enrichment on small-sized biopsies, 5 pairs of forceps biopsies were taken from resected colons of 2 patients. Each pair was isolated with a different soap condition of which 1 sample was isolated with DNAse and the other without. The fold difference between these samples (ΔCt) is plotted. DNAse treatment resulted in a 1.9-fold reduction of human DNA signal (huDNA ratio 0.53, CI: 0.42 −0.65). The bacterial signal was enriched 6.8-fold on average (CI: 2.2-10.52) upon DNA treatment. Triton 0.006% and saponin 0.0125% with DNAse rendered more than 4.3 and 4.5-fold increased bacterial signal respectively in both patients.

Next, we validated our protocol on biopsies from resected colons, which were taken using a forceps to represent clinical biopsies taken during colonoscopy (size: ∼2 mm). 20 biopsies of 2 different patients were taken; patient 1 and 2. Each biopsy was matched with a biopsy that was isolated under similar conditions, but without DNAse treatment. DNAse treatment reduced the human signal in qPCR to 0.53 (CI:0.42-0.65), but increased the bacterial signal 6.8-fold (CI: 2.2-10.52) (figure 3d). Triton 0.006% and saponin 0.0125% gave an enrichment of greater than 4 in both patients (figure 3c). Interestingly, also in absence of soap (PBS control) DNAse treatment resulted in bacterial signal enrichment. This could be explained by the presence of human eDNA due to human cell lysis that may occur during repetitive heating and centrifugation. Ultimately, the bacterial enrichment protocol of strategy 2 was applied in an independent laboratory to isolate bacterial DNA from fish gills. Use of saponin 0.0125% and DNAse treatment doubled the bacterial in qPCR and reduced host signal by factor 135 times, indicating that our enrichment protocol is reproducible and applicable for a wider variety of tissues (see supplementary table 2).

Taken together, our results show that strategy 2, including host cell lysis with 0.0125% saponin and DNAse treatment, successfully decreases human DNA in the sample and boosts the bacterial signal.

### Bacterial composition of human colon tissue by shotgun metagenomics resembles that previously reported by 16S rRNA analysis

Finally, we applied our newly developed approach to clinically acquired colonic biopsies that were isolated usingour optimizedbacterial DNA isolation protocol (supplementary file 1). After degradation of the human DNA, remaining DNA was extracted and analysed with shotgun metagenome sequencing. Metagenomic analysis revealed that the most common phyla were *Firmicutes* (49.5%), *Bacteroidetes* (22.2%), *Actinobacteria* (10.3%), *Proteobacteria* (7.7%), *Verrocumicrobia* (0.6%) and others (9.7%). We compared our data to bacterial composition of human colon tissues reported in literature. Thus far, shotgun metagenomics of microbiomes from tissue samples has been impeded by lack of DNA yield, so shotgun metagenomics has not been reported for colonic biopsies before. Here, we compared our data to samples sequenced by 16S rRNA sequencing (table 1). We found a comparable distribution of bacterial phyla. Furthermore, the Shannon diversity of our study (3.2) was within range of other studies (2.4-3.7). Lastly, our study resulted in an average pairwise UniFraq distance of 0.54 (Fig 5b) which was similar to the UniFraq distance reported in Momozawa *et al*. (0.55). Taken together, with our optimized bacterial DNA isolation protocol (strategy 2) in combination with shotgun metagenomic sequencing, we were able to reproduce previously reported tissue microbial profiles. To our knowledge, this is the first time that colon tissue profiles have been reported with shotgun metagenomics and whereby PCR-induced bias has been omitted.

**Table 1.** Microbiome profiles of human colon biopsies of our study (WGS) resemble those that have been previously published (16S rRNA). We compared our microbiome profiles to those reported in Djuric *et al*., Kiely *et al*., Watt *et al*. and Momozawa *et al*.. These results are represented with a symbol in figure 5. In this table we report the relative abundances of bacterial phyla in percentage. Also, the Shannon index, inverse Simpson index (I. Simpson index) and UniFrac distance (UniFrac d.) are given when reported.

|  | Our study | Djuric *et al.* | Kiely *et al.* | Watt *et al.* | Momozawa *et al.* |
| --- | --- | --- | --- | --- | --- |
| Symbol Fig.5 | Blue star | Red triangle | Red cross | Red hexagon | Red square |
| *Firmicutes* | 49.5 | 61 | 52.5 | 46.5 | -- |
| *Bacteroidetes* | 22.2 | 27.3 | 39 | 43.2 | -- |
| *Actinobacteria* | 10.3 | 2.2 | -- | 0.5 | -- |
| *Proteobacteria* | 7.7 | 4.5 | 2.5 | 5.1 | -- |
| *Verromicrobia* | 0.6 | 3.8 | -- | -- | -- |
| *Fusobacteria* | 0.0 | 0.1 | 1.5 | -- | -- |
| Others | 9.7 | 1.1 | 4.5 | 4.7 | -- |
| Shannon index | 3.2 | 3.5 | 2.4 | 3.7 | -- |
| I.Simpson index | 5.8 | 20.3 | -- | 20 | -- |
| UniFrac d. | 0.54 | -- | -- | -- | 0.55 |

**Figure 5.**
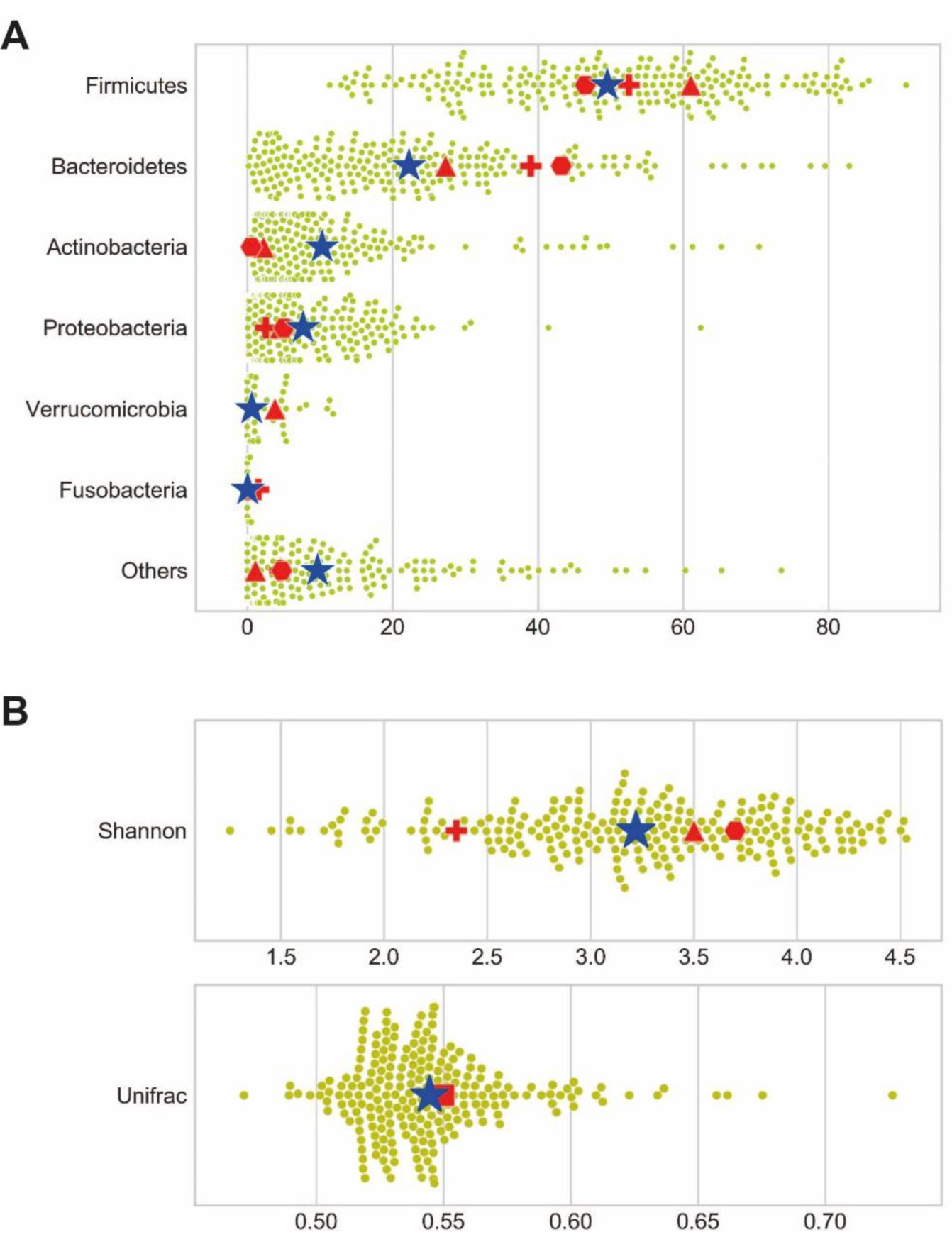
Human colon tissue microbiomes of our study (shotgun metagenomics) versus other studies (16S rRNA). A) The relative abundance of bacterial phyla is shown for study (dots) and the average is marked by a blue star. Averages of Diuric *et al*. (red triangle), Kiely *et al*. (red cross) and Watt *et al*. (red hexagon) are plotted in the graph. The Shannon diversity index and UniFrac distance are represented in B), in which red square represents Momozawa *et al*.

## Discussion

Bacterial DNA isolation from tissues is complicated by large amounts of host DNA. While several strategies, protocols and commercial kits have been developed to tackle this problem, so far none of these considered all elements that we considered important for analysis of tissue microbiomes. In this study we developed a protocol, inspired by Molzym(33), Hasan *et al. (8)*, and the Human microbiome project (HMP) (21), that enriched bacterial DNA through selective lysis of host DNA with 0.0125% saponin and subsequent DNAse treatment. This resulted ina bacterial DNAisolate in which all bacterial subsets were represented, without inducing lysis of bacterial cells or skewing bacterial composition in clinical samples. Of note, our strategy was shown to work also on fish gills and hence can be applied or tailored to other tissues in a similar manner.

We started out testing the Ultra-Deep Microbiome prep-kit (Molzym) in combination with bead-beating (strategy 1), because both methods perform wellin microbiome research (4, 9, 11, 14, 16, 34). The inclusion of bead-beating enhanced isolation of all bacterial phyla, particularly *Actinobacteria* (supplementaryfigure 1). Furthermore, we noticed that the detection of Gram-negative bacteria could be improved by introducing a PBS wash, which we suspect to be caused by the premature lysis of Gram-negative bacteria during the bacterial enrichment steps of this kit (supplementary figure 2). This important limitation has been suggested before (35).

The protocol that we set-up (strategy 2) is an extended version of the protocol that we developed for processing fecal samples (21). This protocol has been modified from the HMP protocol and includes an enzymatic lysis step with mutanolysin, heat-shock and bead-beating. Our bead-beating process has been optimized on a cultured mock community that includes gut bacteria with different susceptibility to lysis. Importantly, fine-tuning of bead-beating speed and duration may be required for each specific bead-beater. It has been questioned whether bead-beating improves bacterial DNA isolation from tissues (36), because it may contribute to some level of DNA degradation (20, 36). However, according to more recent studies, bead-beating does not cause DNA shearing (6, 10) and results in identification of extra species in tissue isolates (18). In our protocol and other studies, bead-beating has proven to result in higher DNA yields (36), more efficient isolation of Gram-positive bacteria (9, 16), a community structure that most closely resembles bacterial input (4), and higher microbial diversity (10). Together, these findings suggest that bead-beating should be included, however it has to be performed with the right type of beads under the right conditions.

Another important stepin our protocol isthe removal of human DNA from the isolate. Previous studies have reported human DNA removal (by qPCR) of roughly >90% in saliva and subgingival plaquesamples with Molysis (15) and >90% in nasopharyncheal aspirate using TurboDNAse(8). Our results showed a reduction of human DNA (by qPCR) of roughly 50% in tissue biopsies. To test whether TurboDNAsewas working well, we tested whether TurboDNAse was able to remove DNA in DNA isolates. These results (not shown) showed that TurboDNAse decreased the DNA concentration by 94%. We conclude that a large amount of human DNA is still inaccessible for DNAse-mediated degradation during our protocol. Interestingly, the use of TurboDNAse without detergent, also increased the bacterial to human DNA ratio. This was also observed before (8). In the study of *Hasan et al*., the use of detergent resulted in a higher pathogen to host DNA ratio, while the attributable effect of detergent was not evident in our study (figure 4c). We suspect that our results are impacted by variety in tissue biopsy size and hence total amount of human DNA. A 2-fold decrease of human DNA signal was associated with an ∼7-fold increase in bacterial DNA signal in qPCR, indicating that human DNA content interferes strongly with the bacterial DNA signal. While it is evident that human DNA remains in the isolate, we have chosen to stick to a mild detergent (saponin 0.0125%) to prevent distortion of the microbiome profile, which may come at cost of complete human cell lysis.

While our protocol is optimized for our research goal, it may require small adaptations for other research objectives. For example, since an important part of our protocol is a DNAse step in which bacterial DNA is still protected by cell wall separation, this DNA isolation protocol may not be optimal to detect bacteria without a cell-wall, like mycoplasma. Study of these types of bacteria requires a different approach, of which antibodymediatedfiltering of bacterial DNA may still be an option. Small adaptations in the protocol may also improve the detection of certain bacterial subtypes, albeit at the cost of less efficient isolation of others. For example, Streptococci DNA-yields may be even higher with more intense bead-beating than in the current protocol. However, we chose to analyze the microbiome as unbiased as possible.

Our shotgun metagenome sequencing results showed that we were able to produce bacterial profiles with Shannon diversity and UniFrac distance that is comparable to 16S rRNA sequencing data of colon tissues, indicating that this sequencing method can be used for tissue microbiome profiling. Nevertheless, small differences were observed between the bacterial composition of our study (shotgun) and three other studies (16S rRNA); we observed fewer *Bacteroidetes* and more *Actinobacteria*. Importantly, similar differences were found in another study comparing shotgun metagenomics with 16S rRNA in stool samples. Ranjan *et al*. reported fewer *Bacteroidetes* with shotgun metagenomics (14-21%) than with 16S rRNA sequencing (34%) and more *Actinobacteria* with shotgun metagenomics (4-7%) than with 16S rRNA sequencing (0.4%) (12). Hence, the differences observed between the colon tissue microbiomes of our and other studies, may be caused by amplification biases.

Taken together, here we showfor the firsttime a protocol to be used for tissue shotgun metagenomics of colon biopsies that omits 16S rRNA amplification steps. Our protocol is mild enough to maintain isolation of Gram-negative bacteria, while it also includes steps that facilitate isolation of sturdy bacteria like *Actinobacteria* and *Firmicutes*. Importantly, our protocol can also be tailored to isolate microbiomes from other tissues, as has been demonstrated by its application to fish gills by an independent laboratory. In other words, our protocol can be immediately used for analysis of stool and colon tissue samples, but may also serve as a foundation for isolation protocols of other study material. Moreover, while we chose shotgun metagenome sequencing, our protocol may also be used in combination with 16S rRNA amplicon sequencing. Thereby our protocol is applicable to many different research settings where it contributes to improved bacterial detection and facilitates analysis of a wide spectrum of bacteria. This way our protocol may contribute to both fundamental and clinical microbiome research, further illuminating the role of microbiome in health and disease.

## Supplementary Data

**Supplementary Table 1.**
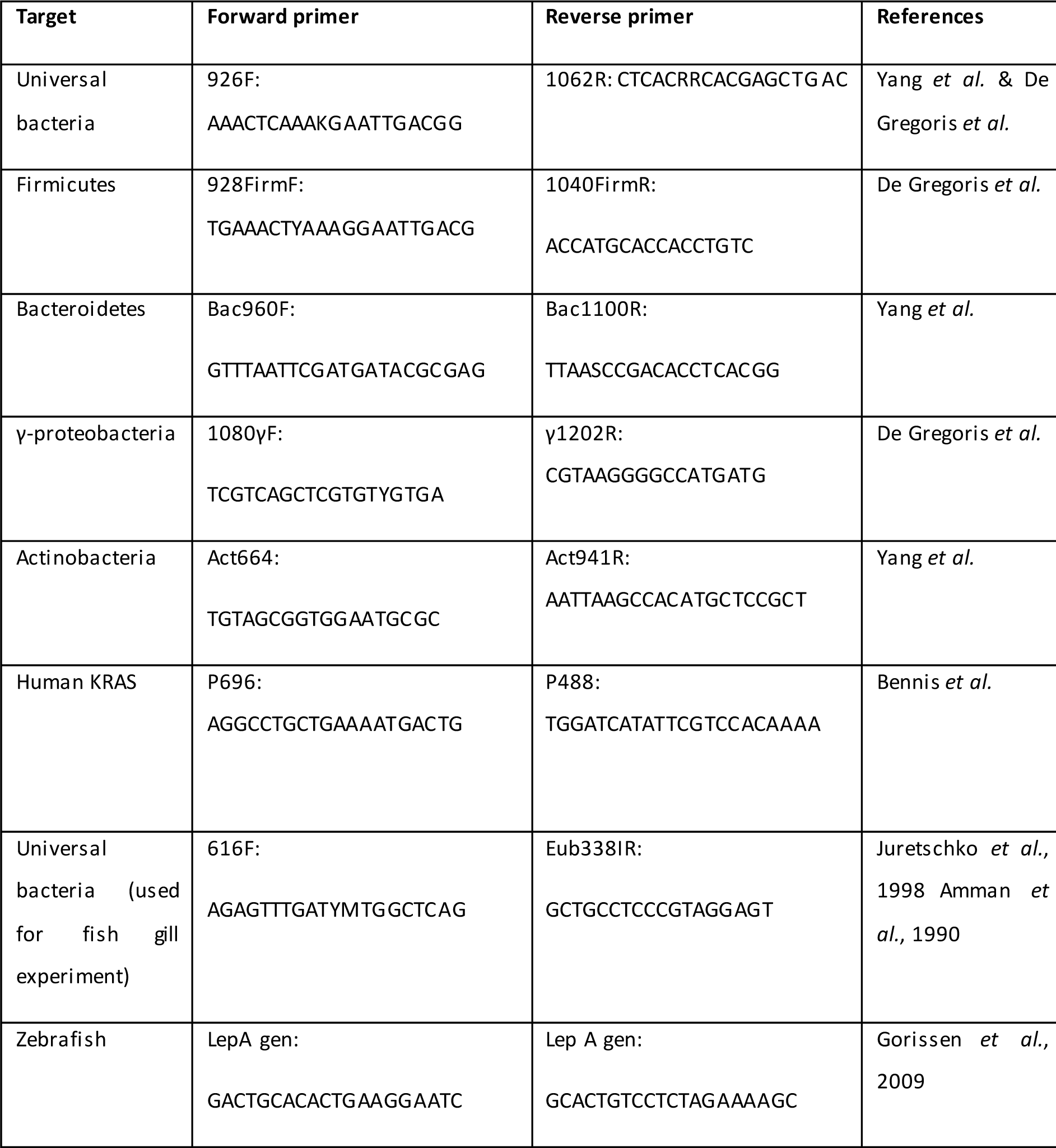
Primers for qPCR.

**Supplementary table 2.**
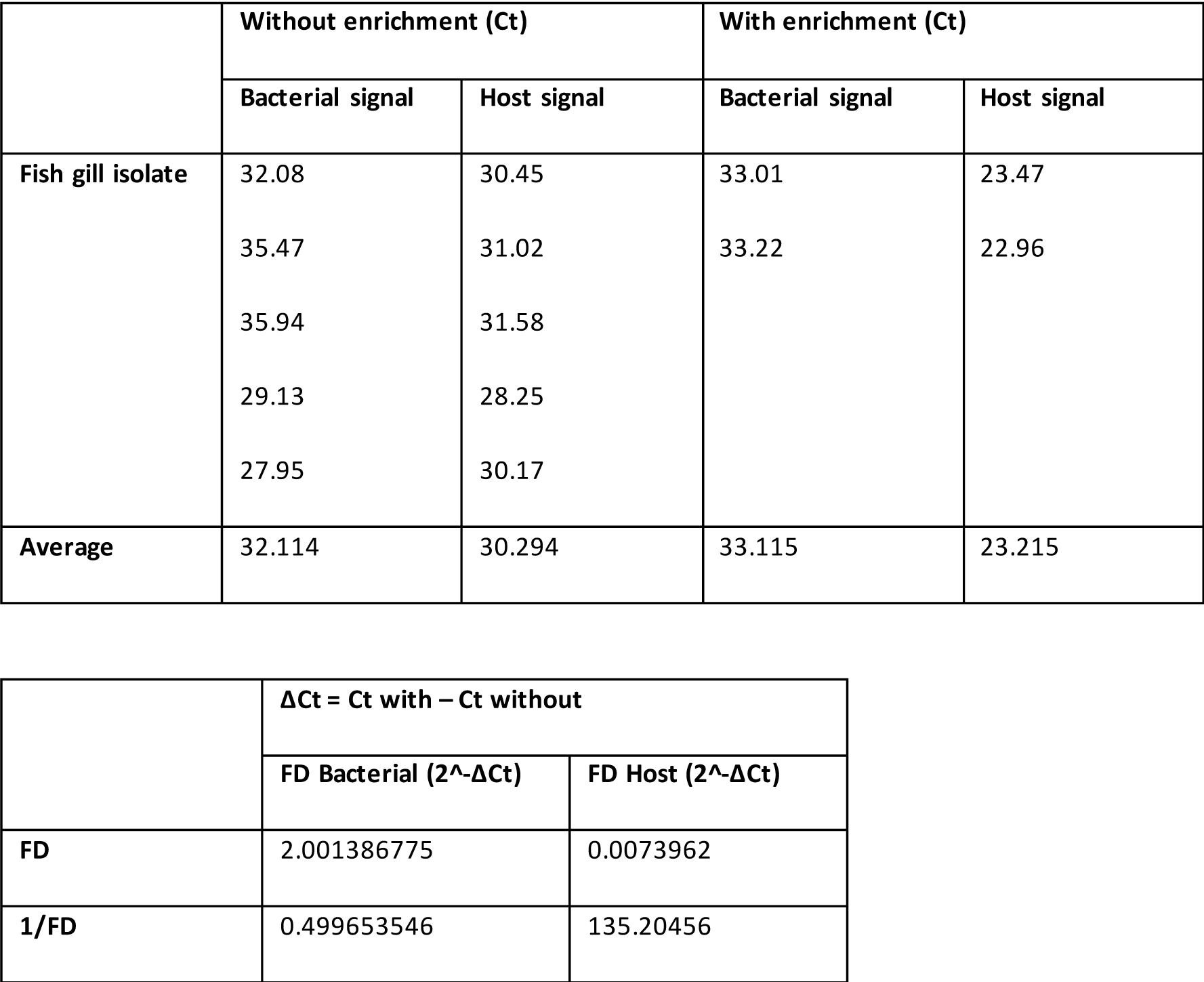
Bacterial enrichment using saponin 0.0125% and TurboDNAse improves bacterial to fish DNA ratio in qPCR. DNA isolations were performed with and without DNAse treatment. Ct values are given in the upper part. In the lower part, the fold difference (FD) between the signal with and without DNA isolation is shown.

**Supplementary figure 1.**
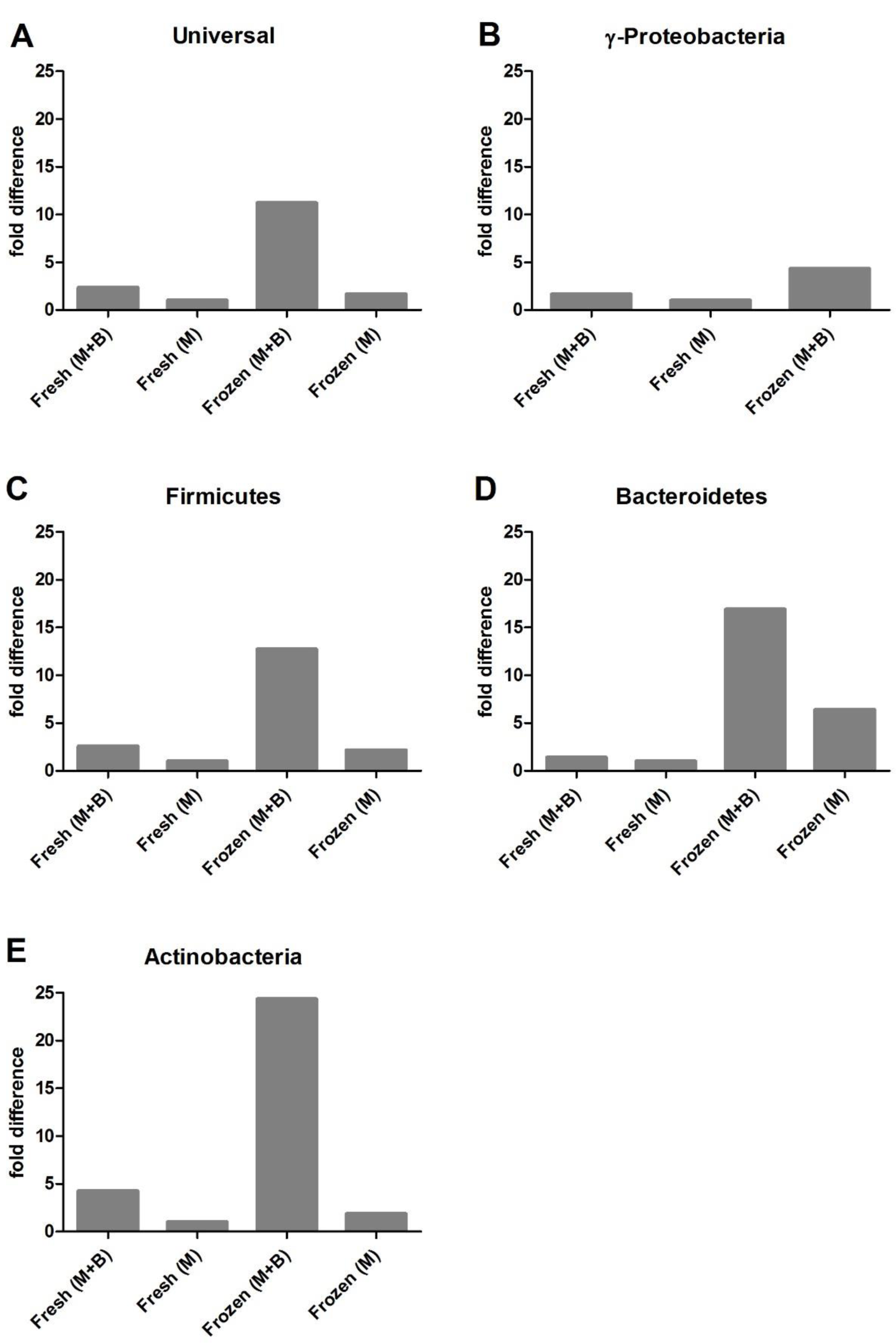
Ultra-deep microbiome prep kit performs better frozen tissue in combination with our optimized bead-beating protocol. Healthy biopsies (∼5 mm) from 1 patient were either snap-frozen (frozen) or immediately isolated with the Ultra-deep microbiome prep kit (fresh). Isolation was either performed with the full protocol provided by Molzym (M) or was combined with bead-beating (M+B). The fold difference represents the bacterial signal relative to the positive control (feces) (ΔCt) and was compared to sample Fresh (M) (ΔΔCt).

**Supplementary figure 2.**
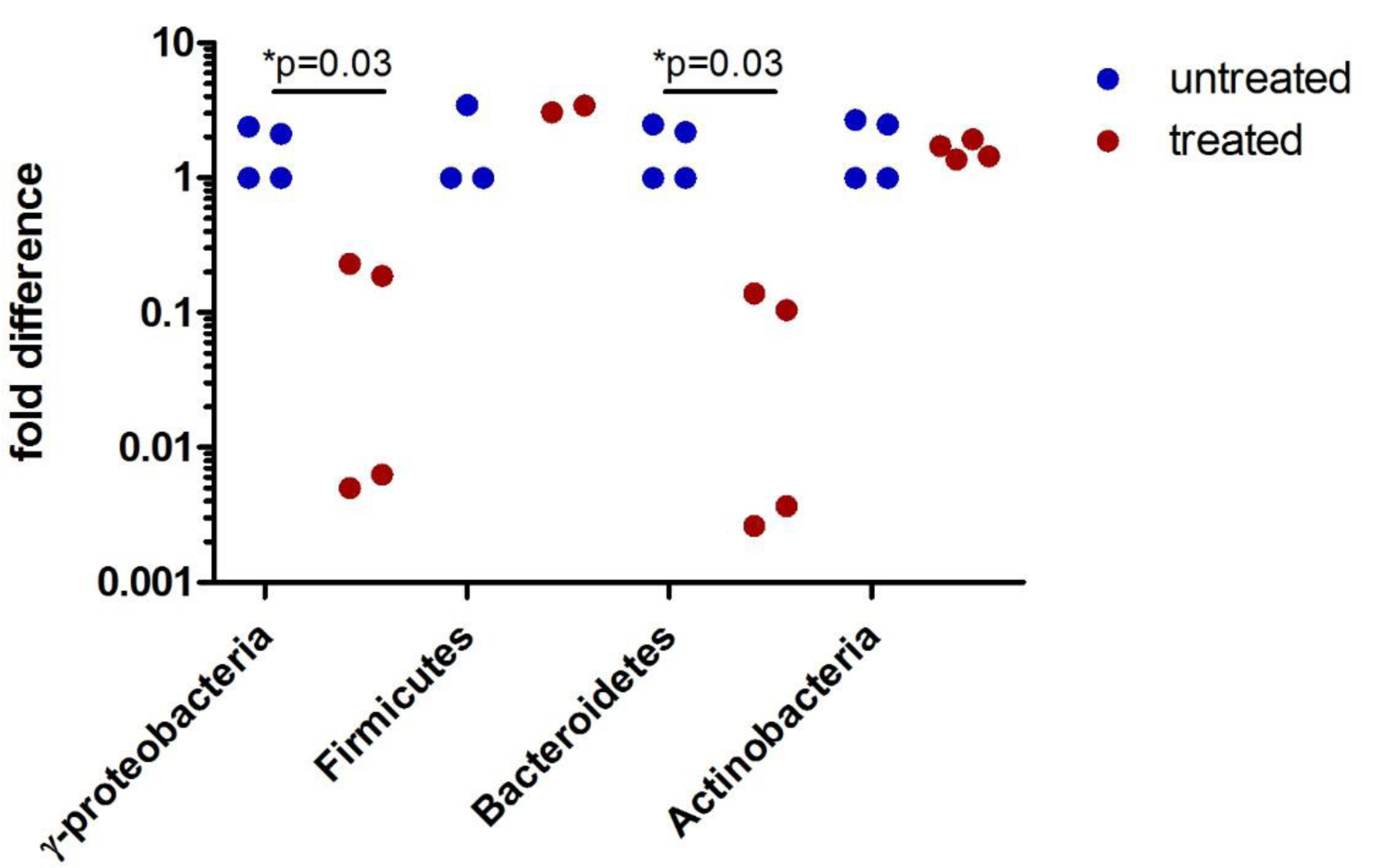
Ultra-Deep Microbiome prep on bacterial mock community results in underrepresentation of γ-Proteobacteria and Bacteroidetes. Two bacterial pellets (mock community) were isolated with the full protocol (treated), whereas 2 pellets were isolated skipping proteinase K, mild lysis and DNA treatment (untreated). To investigate alterations in bacterial composition, each sample was calibrated with its own universal 16s rRNA signal (ΔCt) and was compared to one untreated sample (ΔΔCt). Each sample was run as a PCR duplicate of which both data points were plotted. Mann-Whitney T-test revealed a significant decrease compared to PBS for γ-Proteobacteria and Bacteroidetes.

**Supplementary figure 3.**
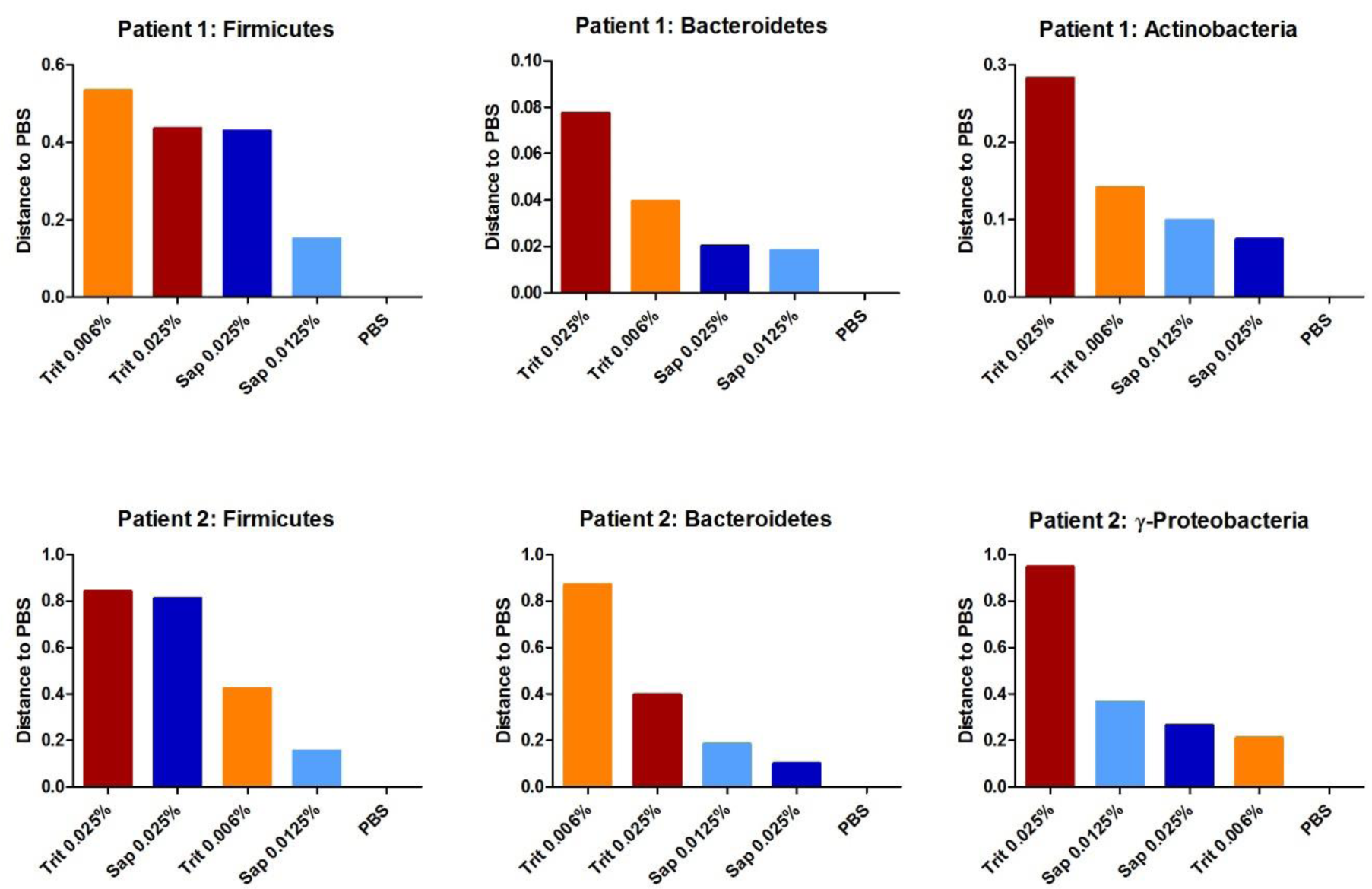
Effect of soap on bacterial composition. Colonic biopsies (∼3 mm) from 2 patients were isolated with our protocol using different soaps and concentrations. The bacterial signal for *Firmicutes, Bacteroidetes, Actinobacteria* and *γ-Proteobacteria* was calibrated with the universal 16S rRNA signal of the same patient (ΔCt) and was compared to PBS sample of the same patient (ΔΔCt). Difference to PBS was plotted.

**Supplementary figure 4.**
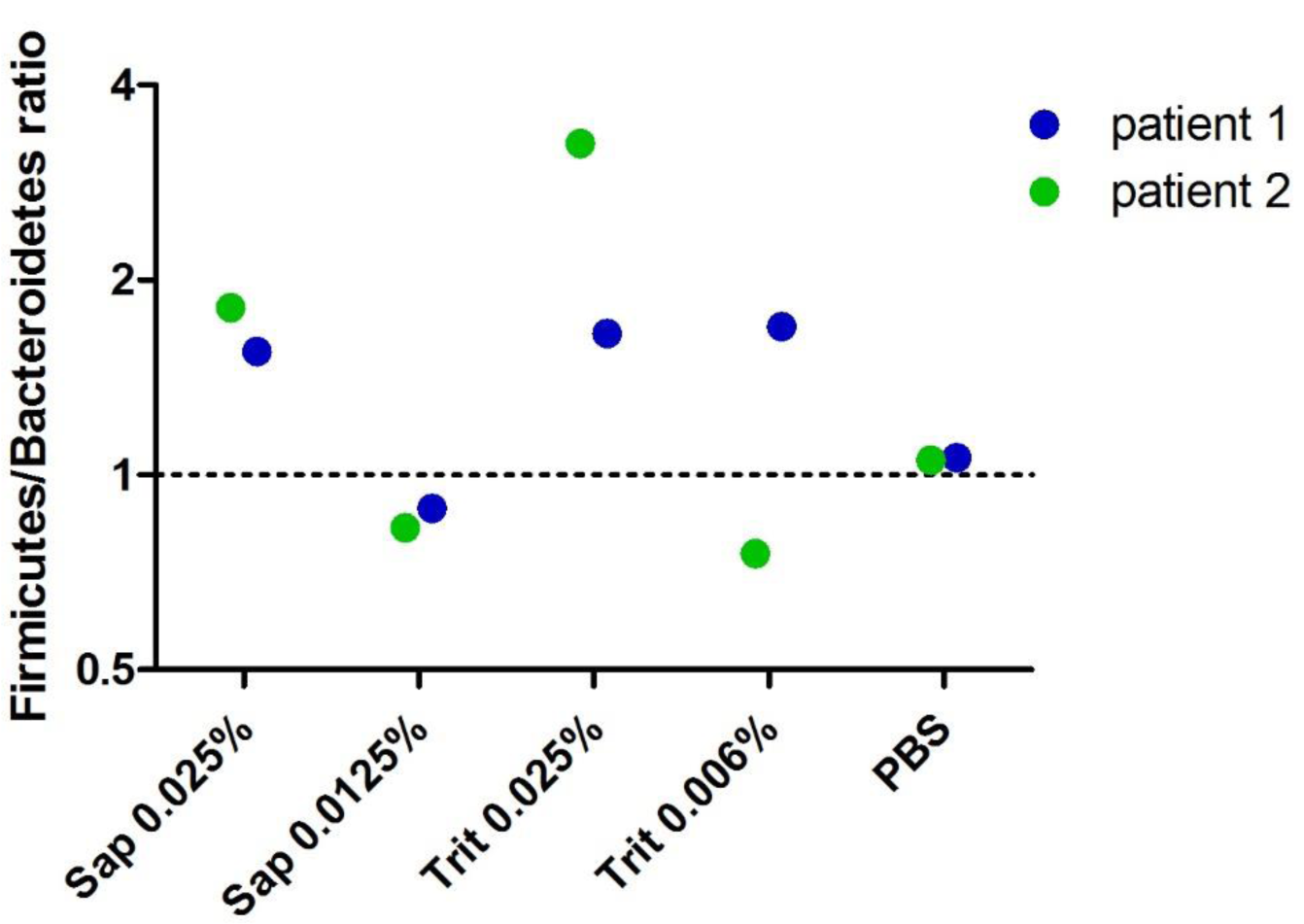
*Firmicutes* to *Bacteroidetes* ratio is least affected by saponin 0.0125%. This graph is extracted from the same experiment as represented in supplementary figure 4. For both *Bacteroidetes* and *Firmicutes* the signal was calibrated with the positive control (mock community) (ΔCt). The enrichment ratio was calculated by 2∧-ΔCt(*Firmicutes*)/2∧-ΔCt(*Bacteroidetes*).

### Supplementary file 1: Protocol

**Bacterial DNA isolation from tissue with bacterial enrichment and bead-beating**.

### Goal

This protocol is optimized for bacterial DNA isolation from human colon tissue samples (∼2 −5mm). During bacterial enrichment, the biopsy is vortexed in PBS to release bacteria from the biopsy. This supernatant (“biopsy wash”) is added back to the sample, after the rest of the biopsy is made into a cell -suspension using proteinase K. The sample is treated with a soap to lyse human cells, which is combined with TurboDNAse treatment to digest external DNA. Subsequently, intact bacteria in the sample are sensitized to lysis using Mutanolysin and heat-shock. Lastly, bead-beating is used for mechanical lysis, which is followed by standard DNA isolation procedures.

Hereby we provide a stepwise protocol, in which blue text represents suggested actions.

### Material

✓ PBS: Tris-HCL (220/12257974/1110, Braun)
✓ Proteinase K (19133, Qiagen)
✓ Saponin 0.0125% (47036-50G, Sigma-Aldrich) in PBS, 0.2µm filtered
✓ TurboDNAse with 10x buffer (AM2239, Qiagen)
✓ Mutanolysin 10 KU in 2mL ddH20 (SAE0092, Sigma Aldrich)
✓ DNeasy Powerlyzer Powersoil kit (Qiagen)
✓ (previously known as MoBio Powerlyzer PowerSoil DNA isolation kit)
  ○ Bead solution
  ○ Solution C1 to C6
  ○ Beads (0.1 mm glass beads)
  ○ 3 sets of 2 mL collection tubes
  ○ 1 set of spin filters

### Preparation

Assure the following:

✓ Clean desk with chloride
✓ Centrifuge at 4°*C*
✓ *70*, 37, 65 and 95 °C *incubator*
✓ Ice bucket
✓ Bead-beater available

### Part 1: Bacterial enrichment

#### PBS wash and host tissue digestion

1. Prepare 2 sets of 1.5 mL Eppendorf tubes, of which 1 set with 500 µL PBS
2. Put frozen biopsies in 500 µL PBS in 1.5 mL tube (use pipettip)
3. Vortex tubes 5 min (speed 8/9) Make PBS/Proteinase K mix
4. Transfer the supernatant (“biopsy wash”) to a new tube and keep on ice
5. If biopsy is ∼2 mm: add 197 µL of PBS and 3 µL of Proteinase K to biopsy For larger biopsies: add 180 µL of PBS and 20 µL of Proteinase K to biopsy
6. Short spin down
7. Incubate samples at 70°C, 400 rpm 15 minutes Set incubator to 37°C
8. Vortex shortly to assist tissue to fall apart
9. Add 700 µL PBS to “biopsy wash” and add to matched biopsy (digested)
10. Spin at 10 000 x g for 10 min 4°C Make Saponin/TurboDNAse/Buffer mix
11. Discard supernatant, save pellet

##### Host cell lysis and DNA digestion

12. Add per biopsy 100 µL mix:
  - 88 µL Saponin
  - 10 µL buffer 10X Turbo DNAse buffer
  - 2 µL TurboDNAse (2 Units/µL)

13. Resuspend by vortexing 15 seconds
14. Short spin down
15. Incubate at 37°C for 30 minutes 400 rpm
16. Add 1.3 mL PBS
17. Centrifuge at 10 000 x g, 10 minutes at 4°C
18. Discard supernatant by pipetting Make mutanolysin mix
19. Add 1 mL PBS and resuspend pellet by vortexing
20. Centrifuge at 10 000 x g, 10 minutes at 4°C
21. Discard supernatant by pipetting
22. Store pellets at −20°C or go to step 23.

#### Part 2: Bead-beating protocol

##### Bead beating preparation

23. Add 180 µL of Bead solution + 20 µL of mutanolysin per sample
24. Resuspend by vortexing
25. Incubate at 37°C for 60 minutes 400 Set up the heater at 65°C
26. ut tubes in the incubator at 400 rpm: 65°C for 10 minutes, heat-up to 95°C (7 minutes) 95°C for 10 minutes
27. Cool down to room temperature and spin down shortly

##### Bead-beating

28. Add 550 µL of Power bead solution to the sample
29. Vortex tubes for 30 to 40 seconds
30. Add mixture to bead-tubes
31. Add 60 µL of solution C1 (first solution of DNeasy isolation kit) Prevent cooling the sample, but bring ice for the following step
32. Bead-beat with the MagNA Lyser:
  - 6400 rpm for 30 seconds
  - On ice for 30 seconds
  - 6400 rpm for 30 seconds Keep samples on ice

##### Bacterial DNA extraction

33. Centrifuge at 10 000 x g for 2 minutes
34. Transfer supernatant to new set of collection tubes *Keep a maximum total volume of 500 µL
35. Add 250 µL of solution C2, Vortex for 5 seconds, incubate on ice for 5 minutes
36. Centrifuge at 10 000 x g for 1 minute
37. Transfer up to 600 - 800 µL to the 2 mL collection tubes
38. Add 200 µL of solution C3, vortex briefly, then place on ice for 5 minutes
39. Centrifuge at 10 000 x g for 1 minute
40. Transfer up to 750 µL of supernatant to the 2 mL collection tubes
41. Add as much as possible without disturbing the pellet (∼850 µL)
42. Shake solution C4, add 1.2 mL (2x 600 µL), Vortex for 5 seconds
43. Add as much as possible, ∼1 mL, avoid that it is so full that it splashes
44. Load approximately 675 µL onto a spin filter, centrifuge at 10 000 x g for 1 minute, Discard the flow (do this 3 until the sample is finished)
45. Add 500 µL of solution C5, centrifuge at 10 000 x g for 30 seconds
46. Discard the flow through
47. Centrifuge at 10 000 x g for 1 minute
48. Carefully place spin filter in new set of collection tubes
49. Add 50 µL of solution C6 to the center of the membrane
50. Centrifuge at 10 000 x g for 30 seconds
51. Discard the Spin Filter
52. Store the extracted DNA at −80°C

### Supplementary file 2: CTAB Extraction

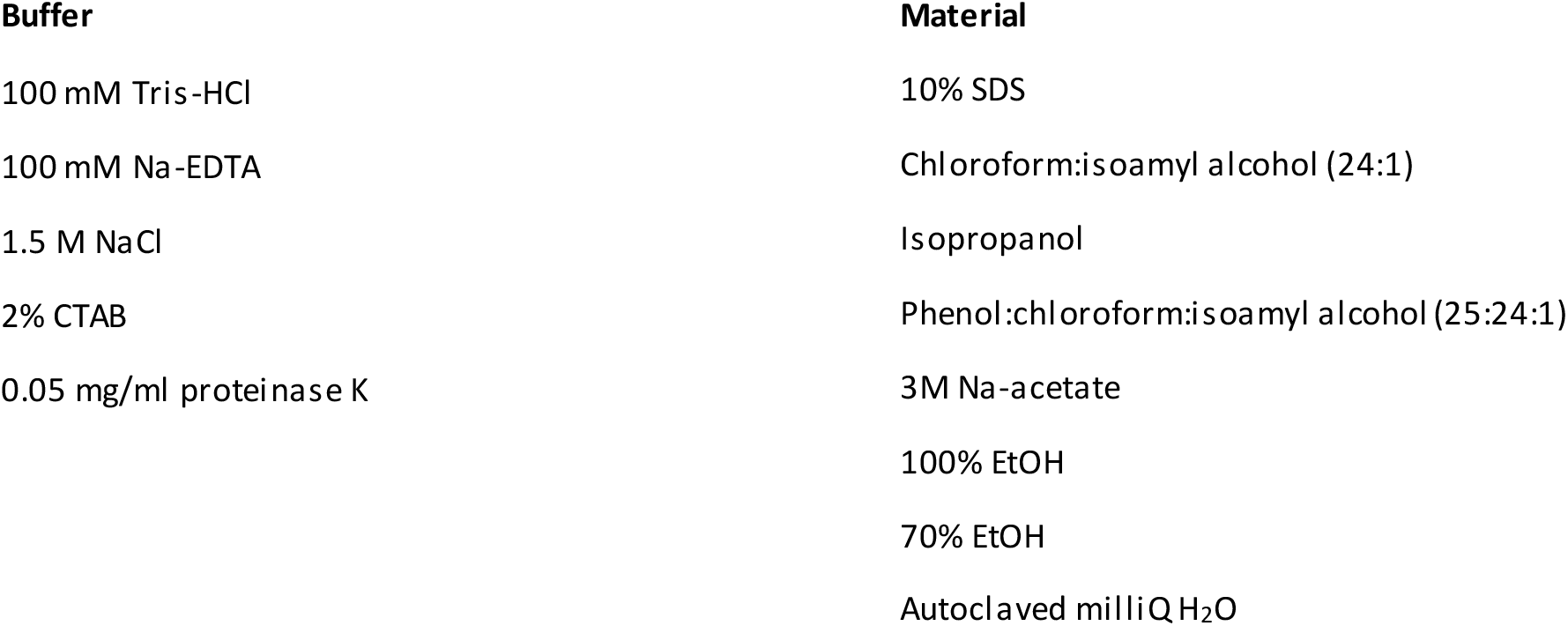

#### CTAB extraction of genomic DNA from de-enriched zebrafish gills

- After the digestion of gill samples with DNase, resuspend washed pellet in 100 µL CTAB extraction buffer and incubate at 37°C for 30 min., mixing every 5 minutes by inverting the tubes
- Add 25 µL 10% SDS to sample, mix well and incubate for 1 hour at 65°C. Mix every 5 minutes by inverting the tubes
- Add 125 µL chloroform:isoamyl alchohol and mix thoroughly for 20 seconds
- Centrifuge samples at max. speed for 15 minutes
- Transfer aqueous phase into clean tubes, discard waste into container in fumehood
- Add 0.6 volumes of isopropanol to samples and incubate overnight at −20°C
- Centrifuge samples at max. speed for 15 minutes
- Pour off isopropanol carefully (don’t lose pellet)
- Wash pellet with 500 µL 70% EtOH, centrifuge 10 min. at maximum *g*
- Pour off ethanol carefully
- Leave tubes open for 5 minutes to evaporate remaining etha nol
- Resuspend pellet in 200 µL autoclaved milliQ

#### RNase treatment of DNA extractions

- Add 1 µL (10 mg/ml) RNase A to samples, incubate at 37°C for 30 minutes
- Add 200 µL phenol:chloroform:isoamyl alcohol, mix thoroughly for 20 seconds
- Centrifuge 15 min. at maximum speed
- Transfer aqueous phase into new tube, discard phenol waste into container in fumehood
- Add 2 volumes of 100% EtOH and 0.1 volume of NaAc, mix by inverting tube
- Incubate at −20°C for 1 hour
- Pellet DNA by centrifuging for 20 minutes at max. speed
- Wash pellet with 500 µL 70% EtOH, centrifuge 10 minutes at max. speed
- Pour of ethanol carefully, spin down the rest of the ethanol by short centrifugation
- Remove residual ethanol by pipetting, without disturbing the pellet
- Dry pellet until all ethanol is evaporated
- Resuspend pellet in 50µL autoclaved milliQ water

### PCR

#### qPCR programme

|  |  |  |
| --- | --- | --- |
| 3:00 | 96°C | 1x |
| 0:15 | 96°C | 40x |
| 0:20 | 58°C |  |
| 0:30 | 72°C |  |
| 2:00 | 72°C | 1x |

#### qPCR mix

|  |  |
| --- | --- |
| SYBR mix 2x | 10 µL |
| Forward | 0.6 µL |
| Reverse | 0.6µL |
| H2O | ... µL (upto 20 µL) |
| DNA | 5 ng |

